# A minimal 3D model of mosquito flight behavior around the human baited bed net

**DOI:** 10.1101/2020.01.21.913608

**Authors:** Jeff Jones, Greg Murray, Philip J McCall

## Abstract

Advances in digitized video-tracking and behavioral analysis have enabled accurate recording and quantification of mosquito flight and host-seeking behaviors, enabling development of Individual (agent) Based Models at much finer spatial scales than previously possible. We used such quantified behavioral parameters to create a novel virtual testing model, capable of accurately simulating indoor flight behavior by a virtual population of host-seeking mosquitoes as it interacts with and responds to simulated stimuli from a human-occupied bed net. We describe the model, including base mosquito behavior, state transitions, environmental representation and host stimulus representation. In the absence of a bed net and human bait, flight distribution of the model population is relatively uniform in the arena. Introducing an unbaited net induces a change in distribution due to landing events on the net surface, predominantly occurring on the sides and edges of the net. Presence of simulated human baited net strongly impacted flight distribution patterns, exploratory foraging, the number and distribution of net landing sites, depending on the bait orientation. As recorded in live mosquito experiments, contact with baited nets (a measure of exposure to the lethal insecticide) occurred predominantly on the top surface of the net. Number of net contacts and height of contacts decreased with increasing attractant dispersal noise. Results generated by the model are an accurate representation of actual mosquito behavior recorded at and around a human-occupied bed net in untreated and insecticide treated nets. In addition to providing insights into host-seeking behavior of endophilic vectors, this fine-grained model is highly flexible and has significant potential for in silico screening of novel bed net designs, accelerating the deployment of new and more effective tools for protecting against malaria in sub-Saharan Africa.

## Introduction

Whether to combat insecticide resistance, exploit new knowledge of vector biology, increase community acceptance or to accommodate changes in abiotic conditions, mosquito vector control methods are under constant pressure to improve. In sub-Saharan Africa, where 90% of malaria occurs, Long Lasting Insecticidal Nets (LLINs) have been the main driving force in the reduction of malaria cases [1], [2], but widespread and growing insecticide resistance to the pyrethroid insecticides used on nets threatens to stall or even reverse recent advances [3], [4], [5], [6]. Development of novel insecticide compounds for use either alone or in combination with existing pyrethroids, provides one solution [7], [8] but raises significant questions regarding the optimal placement of compounds on the net walls or roof, both for maximizing efficacy and minimizing cost. A detailed understanding of the spatio-temporal nature of mosquito responses to various insecticidal treatment(s) on the bed net interface is critical to balance these competing requirements for the numerous combinations possible.

The development and eventual implementation of a new bed net is an expensive and time-consuming process and the pipeline from early phase screening of chemicals with potential through the range of laboratory and field tests needed to generate the evidence required by regulatory authorities before the commodity finally reaches the affected communities, can take up to ten years.

Recent technical advances in tracking and analysis of mosquito flight behavior during host seeking [9], [10] have enabled the detailed characterization of flight patterns at the human host, ultimately revealing how insecticide treatments on bed nets affect the behavior of malaria vector mosquitoes. The growing body of data arising from those studies not only builds the evidence base required to accelerate the development process, but it also provides an excellent foundation for developing models of host-seeking behavior and experimentally validate the new tools at earlier stages. This paper describes one such model: a fine-grained agent-based approach for modelling how indoor insecticide treatments deployed as residues on bed nets affect the behavior and survival of mosquitoes.

To capture the complex interactions between mosquitoes and a human host lying beneath an insecticide-treated bed net, individual mosquito flight paths and local interactions within the indoor environment are required. Yet most modelling approaches to disease transmission by insect vectors set the scale at a much higher level, emphasizing interactions at village scale (and above) [11], [12]. Such large-scale models are informed by a substantial body of similarly large-scale epidemiological datasets e.g. morbidity or mortality within local medical centers, ultimately aggregated at national level.

At the opposite end of the scale there is a substantial body of experimentally generated data on the behavior of individual mosquitoes in experimentally controlled settings such as wind tunnels and olfactometers [13]. A wide range of possible sensory modalities could influence host-seeking selection behavior in these conditions and these different potential stimuli have been investigated with regard to isolated CO_2_ concentration, olfactory [14], [15], [16], visual [17], [18], auditory [19], [20] and tactile sensory cues [21], or combinations thereof [22].

There is a knowledge gap between individual mosquito behavior and downstream effects in terms of epidemiological disease transmission that could be filled by fine grained models of insect flight behavior and host interactions. However, this is hampered by uncertainty regarding the relative importance of the different sensory modalities which mosquitoes utilize during host-seeking. Agent-based modelling is a useful approach because it can represent stochasticity and heterogeneity at fine-grained scales and generate emergent behaviors not explicitly encoded into the model [23]. Previous agent-based approaches have included a 2D model of insect flight to study orientation and tracking of odor plumes [24], an agent-based model of the effect of different urban environmental factors on population distribution [25]. A 2D model of mosquito interactions with LLINs was used to assess community-scale protection [26]. The influence of *human* movement on malaria transmission was studied with an agent-based model in [27] who found that the spatial locations of malaria hotspots was strongly influenced by human movement.

In this paper we propose to circumvent the debate regarding the relative importance of particular sensory stimuli by developing a minimally simple agent-based model of 3D mosquito flight behavior. The model is stimulus-agnostic and uses a single generic ‘attractant’ signal emanating and dispersing from a host. We use the model to study mosquito flight behavior and landing distribution patterns on unoccupied and host-occupied bed nets. By validating the parameters of the model with previous experimental results, we hope to apply the model for use as a virtual experimental tool for predictive vector control studies.

Initial studies on mosquito behavior at bed nets used adhesive coated netting to map the final position of mosquitoes at the bed net surface; by showing a preference for the net roof, the attractant plume rising from the human host beneath was described indirectly [28,29]. This approach is limited as the initial contact site might not be the ultimate preferred destination while mosquitoes adhering to the net surface are essentially removed from the ‘pool’ of mosquitoes available to study further activity.

Recent technological developments in imaging and computing have enabled accurate tracking of mosquito flight dynamics. This is achieved by analyzing video recordings of mosquitoes and applying image processing and analysis methods to reconstruct and log the flight paths and range from relatively simple approaches using a single camera [30], [31] to more complex multi-camera methods [9], and 3D reconstruction approaches [32], [33]. These multi-camera approaches allow the tracking of mosquito flight around a bed net and therefore a more complex representation of bed net occupancy at a ‘life-size scale.

In [10], the peripheral regions surrounding the net were subdivided into polygonal regions, allowing the differential distribution of mosquito activity to be recorded over time and under different bed net occupancy conditions. Data of this type may be used to inform models of mosquito flight and behavior, with the aim of testing variations of vector control tools in-silico.

## Materials and methods

InVeCTS (Indoor Vector Control Testing System) creates a virtual environment in which to assess mosquito populations’ interactions with their host and environment. It uses an agent-based approach with fine-grained spatial representation in which a mosquito population can interact with a human host emanating the hypothetical spatially distributed attractant stimulus over time. Mosquito flight occurs in real time and all mosquito flight paths and interactions with the environment are recorded for subsequent analysis.

### Environment and Mosquito Representation

A population of 25 mobile virtual mosquitoes is created. These individuals fly in a continuous 3D space representation inside a discretized spatial arena, representing an insectary whose dimensions directly correspond to an experimental insectary at LSTM (5.6m long x 3.6m wide x 2.3m high). This virtual insectary can contain a bed net and human host, as shown in Fig. 1.

**Fig 1.**
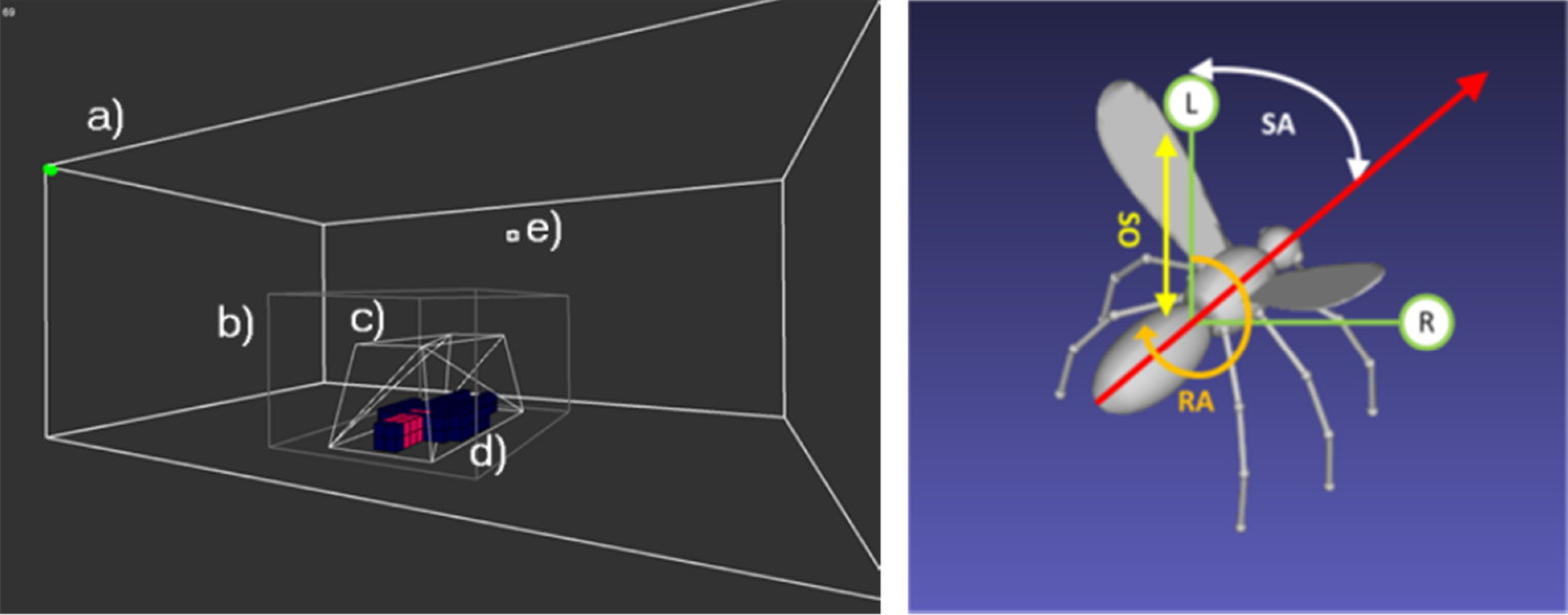
Arena and mosquito representation in the InVeCTS model. Left: Mosquito flight environment. The environment consists of the main insectary arena comprising (a) a cuboidal recording volume corresponding to that used in [10] (b), a bed net (c) containing a human host (d). Mosquito release location is indicated in (e). Right: Schematic representation of virtual mosquito agent with its current directional vector (long arrow), its offset Left and Right sensors (L,R), Sensor Offset distance (SO), Sensor Angle (SA) and Rotation Angle (RA) parameters.

The population is introduced at the release site (Fig. 1e) and begins exploration of the arena. A hypothetical attractant plume is projected with the size and shape of a human host. To represent and store the attractant profile the arena is divided into a 56 × 36 × 23 cubic lattice of 10cm^3^ cells. The host bait profile is configurable to represent hosts of different sizes (Fig. 1d) and the cells making up the bait profile are color-coded to indicate regions where greater concentrations of attractant are emanated. As previously stated, a wide range of environmental cues (for example, *CO_2_,* skin odors, sound, vision, temperature) are known to influence mosquito behavior but their relative contribution and sequential importance in host location is still uncertain ([34], [35], [36]). To circumvent this uncertainty, and to simplify the model, we utilize a generic representation of a single spatial attractant, emanating and diffusing from the host. We use a simplifying assumption that the multi-modal nature of the attractant profile can be represented in this manner. We use a very simple cellular automata-based dispersion mechanism in order to enable real-time dispersal of the attractant profile.

### Projection and Dispersal of Human Bait Attractant

At each scheduler step attractant is projected at bait profile locations and isotropic dispersion is implemented in parallel for all cells in the 3D lattice volume by distributing the attractant between the current cell and the 26 nearest neighbor cells. Vertical dispersion (a very coarse approximation of convection) is implemented for all cells by dispersing a fraction of each cell to the cell immediately above the current cell in the arena. We use such a simple scheme for two reasons. Firstly this is because of the uncertainty of the relative importance of different environmental cues, and secondly, because we wish to visualize the dispersal in real-time which is not yet possible with more complex representations of environmental dispersion.

Real-time projection, diffusion and convection of bait profile attractant in the lattice is parameterized by independent weight parameters for diffusion (*W_d_*) and convection (*W_c_*). At every scheduler step a generic chemo-attractant, weighted by the bait profile map (to account for higher regions of attractant, for example: from the mouth) is projected into the 3D attractant lattice at regions directly corresponding to the bait map profile shape.

#### Attractant Plume Diffusion

At every scheduler step, volumetric diffusion of attractant is approximated by the following cellular automaton-based method.

For each cell in attractant lattice (axes x,y,z): *C_x_yzt* + 1 = ((*C_x_yz* + local neighborhood attractant values (radius 1))/27) * Wd

To approximate noisy dispersal of attractant (for example, turbulent air currents) and their effects on host-seeking behavior each cell can be modified by adding a value randomly sampled from a Gaussian distribution with mean 0 and standard deviation 1. Noisy dispersal can be increased by multiplying this noise value by a scaling parameter *D_s_*.

#### Attractant Plume Convection

At every scheduler step volumetric convection upwards is approximated by copying of vertically stacked horizontal ‘sheets’ of cells containing attractant values to cells in the sheets above (Insectary ceiling is set to *Y* = 0, layers below are *Y* + 1 etc). Projection is weighted by convection weight parameter *W_c_*.

For each cell in attractant lattice (axes *x, y, z*): *C_x_yz t* + 1 = (*C_x_y* + 1*z*) * *W_c_*

A visualization of the effect of attractant plume dispersal is shown in Fig. 2. We acknowledge and reiterate that this is a very simple approximation of attractant dispersion. However, it is sufficient to generate a stable 3D attractant field and avoids emphasising any one particular (potential erroneous) stimulus type over others.

**Fig 2.**
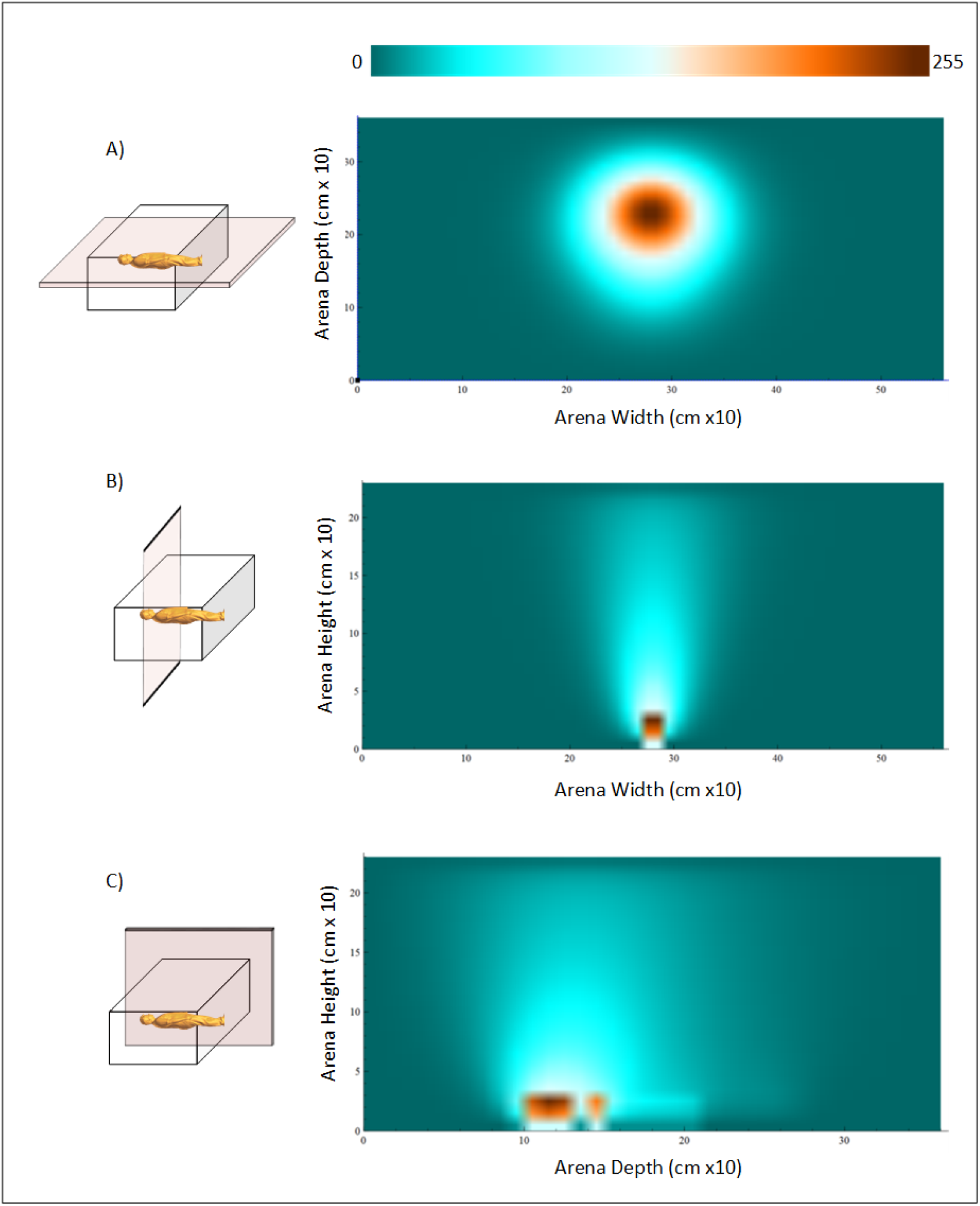
Visualization of attractant plume within the arena. Left side shows orientation of cross-sectional slice through attractant field within the arena. Right side shows attractant concentration scaled individually for each image to the look-up table (top). a) Gradient concentration of attractant plume (looking from above insectary, slice taken from coronal plane, 0.7m above floor) b) Gradient concentration of attractant plume (looking along axial plane of human profile). Section is taken at the head region of the human profile. c) Gradient concentration of attractant plume (looking at side-on human profile). Section is taken as a saggital slice along the middle of the human profile.

### Mosquito Sensory-Motor Algorithm

At each scheduler step the mosquito samples the attractant value in the insectary from two offset sensors (sensor offset distance may be adjusted with the parameter **SO**. Sensor angle may be adjusted with the parameter **SA**. If the sensor values are different the agent orients itself locally in space by rotating about its own axis towards the strongest value, by the amount in degrees given by the Rotation Angle (**RA**) parameter. Additionally, the agent orientation is subject to random modification by the **pCD** (probability of Change Direction) parameter. If a randomly sampled value from the uniform distribution is < **pCD** (default value 0.25), the agent will select a random orientation in 3D space. At each scheduler step each agent moves forwards in its current orientation by a velocity given by the parameter **V** (default value 1cm).

**Table 1.** Model parameters. Parameters are divided into groups reflecting the effect on the environment, agent sensory-motor behavior, and agent behavioral transitions.

| Parameter Type | Name | Default Value | Description |
| --- | --- | --- | --- |
| Environment | Wd | 0.7 | Diffusion damping factor |
|  | Wc | 0.3 | Convection damping factor |
|  | Ds | 0 | Dispersal noise value |
| Mosquito Flight | SA | 45 | Sensor angle (deg) |
|  | RA | 22 | Rotation angle (deg) |
|  | SO | 1 | Sensor offset distance (cm) |
|  | V | 1 | Velocity (cm) per scheduler step |
| Mosquito State | pCD | 0.25 | Probability of changing direction |
|  | pRest | 0.1 | Probability of resting on surface |
|  | pLeave | 0.05 | Probability of leaving surface |

### Mosquito behavior Transition Function

Agent behavior in response to environmental cues is determined by a set of states and the subsequent transition between these states. To avoid a fully deterministic response to population behavior, transition between states is mediated by probabilistic sampling, implemented by individual parameters, corresponding to a Markov process of state transitions (see Fig. 1 for all model parameters). A schematic overview of the behavioral transitions can be found in supplement S3. The agents are initially in the **PRE-RELEASE** state and their state is updated every scheduler step. A mosquito enters the **FLYING** state when a randomly sampled value from a uniform distribution is < 0.1. If an agent is at a wall, ceiling, bed net, or floor surface it will rest if a randomly sampled value from a uniform distribution is < **pRest** parameter. Non-resting mosquitoes change direction to a random orientation and resting agents stay in the current location until a randomly sampled value from a normal distribution is < **pLeave** parameter. It the agent lands on, or is resting on a treated bed net its starting health value (100) is decremented by a toxicity value *t*. If the health of an agent reaches zero it is removed from the arena.

### Parameter selection and model validation

Model parameters were set to match values obtained by previous image tracking experiments in the same LSTM insectary space. Dimensions of the virtual arena were identical to the insectary, as were positions of the bed net, host position, number of mosquitoes, mosquito release location, and experimental run time. Although measurements of mosquito flight speed vary in the literature (and indeed during different mosquito behaviors) a mosquito flight speed of 300*mms* was selected to approximate the flight speed found in the tracking studies in [9].

Varying the *SA* and *RA* parameters affects flight tortuosity which subsequently affects arena occupancy and foraging behavior. After evaluating a range of *SA* and *RA* angle combinations, we settled on fixed *SA* of 45° and a *RA* angle of 22° (see supplement S6 for evaluation and description of tortuosity calculation). This resulted in path tortuosity which approximated those of the image tracking experiments reported in [10]. Limiting assumptions of the model are described in the discussion section.

### Model scheduler and model experiment setup

At the start of an experiment, the system creates the virtual insectary environment and the agent population. The population is placed at the release location and the model begins iterating through its main run-time loop as shown in Fig. 3.

**Fig 3.**
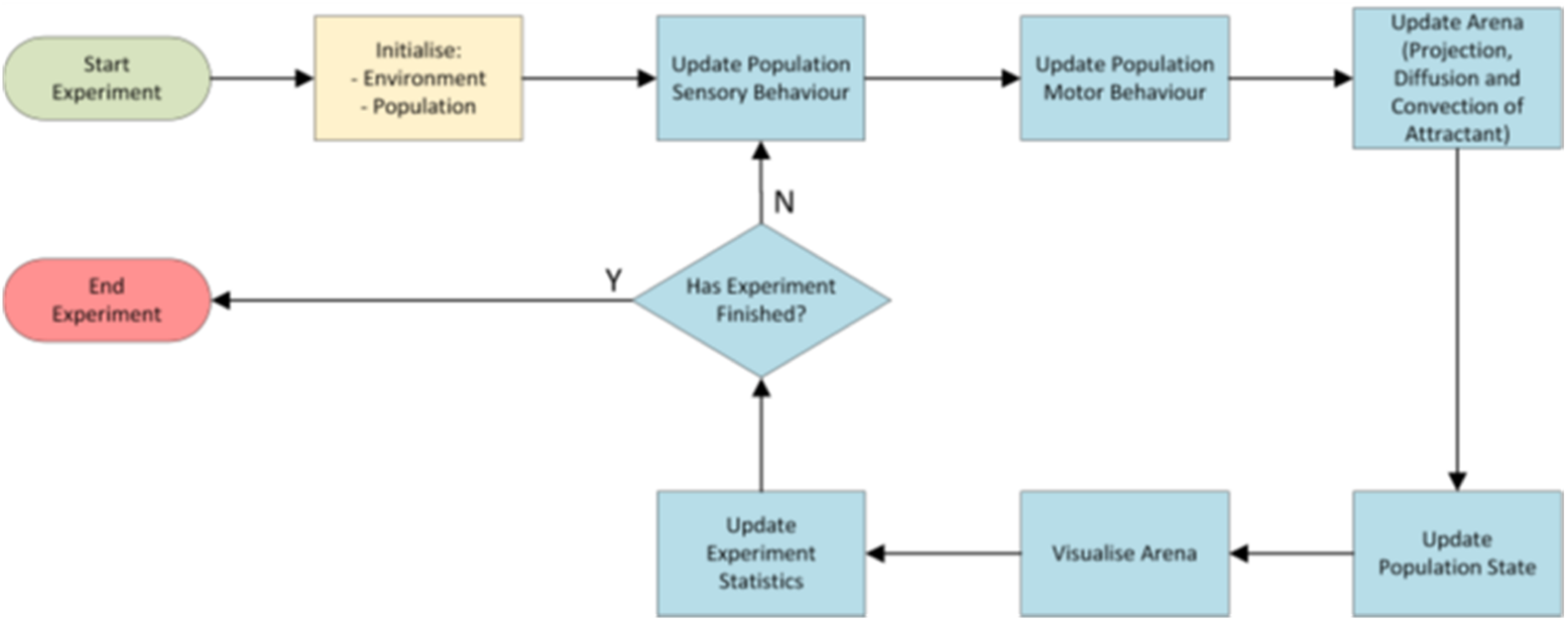
InVECTS model scheduler.

Each scheduler step comprises 1/30*th* second. The model halts after 1 hour, or 108000 scheduler steps. After the model halts, log files pertaining to virtual mosquito activity (three-dimensional coordinate positions, agent state and agent health), bed net contact locations, and agent presence within spatial regions surrounding the bed net are saved to disk for further analysis.

The user can interact with the model during run-time, visualizing the current state of the model and population flight behavior in 3D space using a mouse. Specific experimental configurations can be saved and re-loaded for repeated runs.

Five control experiments were performed in the virtual insectary where no bait or no bed net was present, as a baseline for the flight behavior of the model. Twenty experiments were performed for each of the human bait conditions (no bait, left oriented bait, and right oriented bait). To assess the effect of noisy attractant dispersal we the used the left oriented bait profile and performed five experiments at each noise parameter value.

## Results

### Spatio-temporal flight activity and foraging behavior

Control experiments, where neither bait nor bed net were present, established baseline flight behavior and distribution. Occupancy of all regions is plotted in Fig. 4 across the X and Z planes. The heatmap images indicate a top-down summary of occupancy from above the entire insectary results, demonstrating that in the absence of a host or bed net, flight activity was relatively uniform throughout the arena (Fig. 4,A).

**Fig 4.**
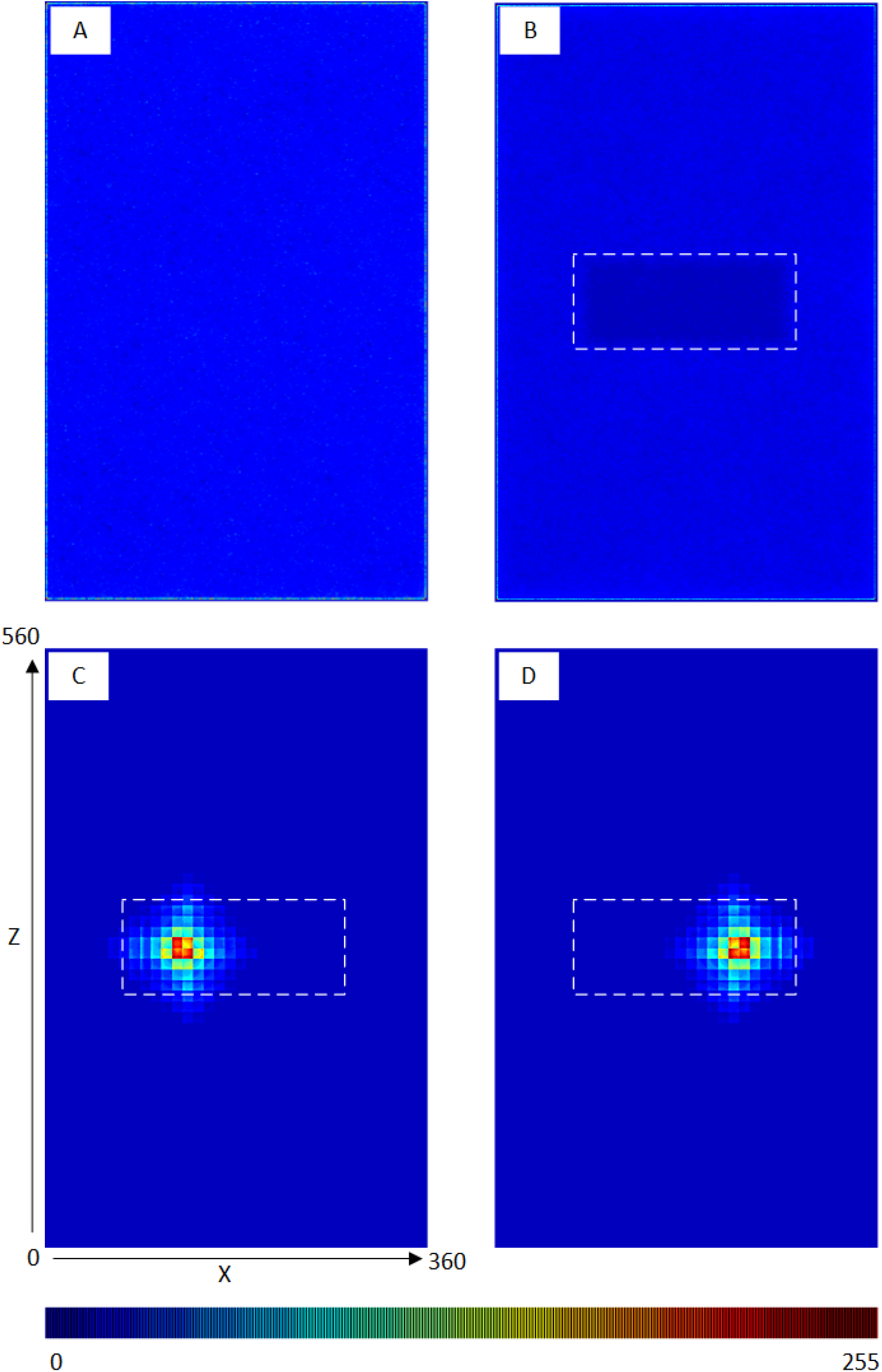
Flight distribution of virtual mosquito population in the virtual insectary. Distribution of activity in the XZ plane (i.e. looking from above). Bed net position indicated by dashed rectangle. A) Insectary containing no bait and no bed net, B) Insectary with unbaited bed net, C and D) Insectary with left and right oriented human bait. Occupancy at all heights binned to 1 pixel range and scaled to 8-bit look-up table color values (bottom).

When an unbaited net is introduced to the arena, the flight distribution is still relatively uniform except in those regions containing the empty bed net where the mosquito population cannot enter (Fig. 4,B). When the stimulus profile of a human shape is placed on the bed, within the net, the flight distribution shows a marked change, with the population showing a strong preference to fly in the regions of the strongest diffusing and convective stimuli, corresponding to the orientation of the human bait profile (Fig. 4,C and D).

As in nature ([10]), in the InVeCTS model, foraging by the virtual mosquito population is strongly affected by the presence or absence of a human host. Fig. 5 (top) demonstrates spatial foraging. In the presence of bait, a narrow field of the arena is explored, whereas the virtual population explores a much larger area of the arena when no host is present. Temporal exploration of arena occupancy over time (the total fraction of the arena explored by the population) confirms that the presence of bait results in less foraging (an exploitation strategy). When bait is absent the population adopts an exploration strategy. This behavioral response is induced by the presence of dispersed attractant stimuli within the arena. Example video recordings of short model runs in unbaited and baited conditions are shown in S1 and S2 respectively.

**Fig 5.**
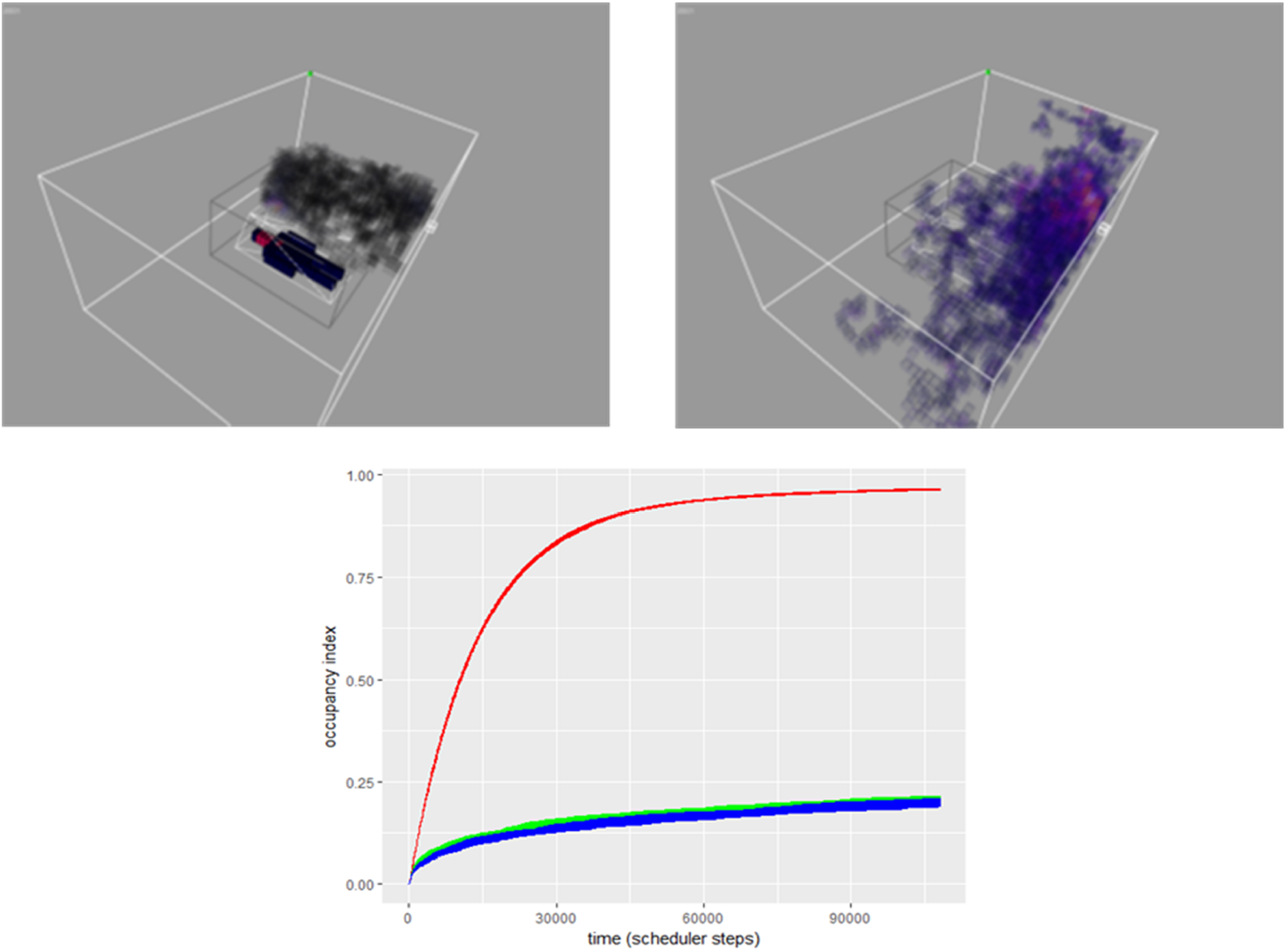
Effect of host presence on foraging and arena occupancy. top: Snapshot of occupancy in a baited (left) and unbaited (right) arena after 20,000 scheduler steps. Voxels indicate regions of the virtual insectary which have been occupied. bottom: Occupancy index (the fraction of the arena explored by the population) for an unbaited bed net (red) compared to baited conditions (green represents left sided bait and blue is right sided bait).

### Flight Path Tortuosity

Flight path tortuosity was calculated using the rolling tortuosity metric described in S6. A sliding window of 50 mosquito movements along the flight path was assessed for all points and only uninterrupted segments of flight paths were used (i.e. those not including landing and resting). Path tortuosity under different bait conditions are shown in Table 2. Tortuosity in unbaited arenas was slightly lower than measured previously in [10], however that study only included information from the recording volume captured by the cameras (where tortuosity is increased by the mosquito responses to the net). The model tortuosity includes the entire insectary space with more free flight space which reduces overall tortuosity. Under baited conditions (which attracts greater mosquito activity at the net) the model tortuosity closely matches experimental values.

**Table 2.**
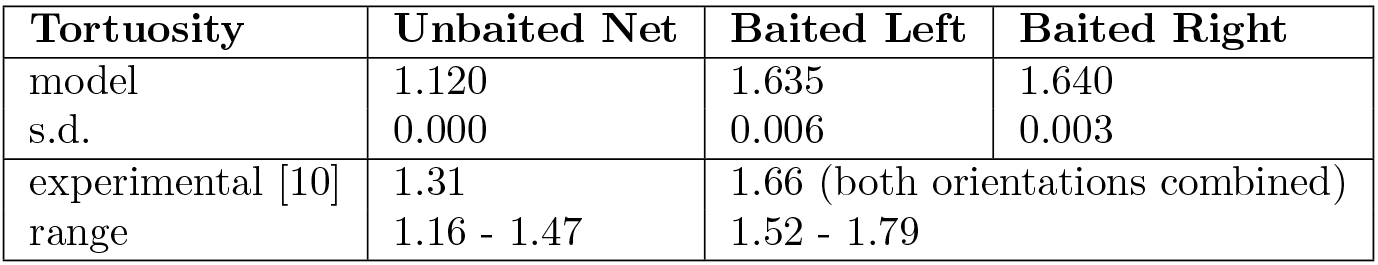
model vs experimental flight tortuosity

### Net contact distribution

Net contact locations in the unbaited condition occurred mainly along the sides of the bed net, particularly along the longest axis of the net (Fig. 6, top). There was also relatively little contact at the top surface of the net.

**Fig 6.**
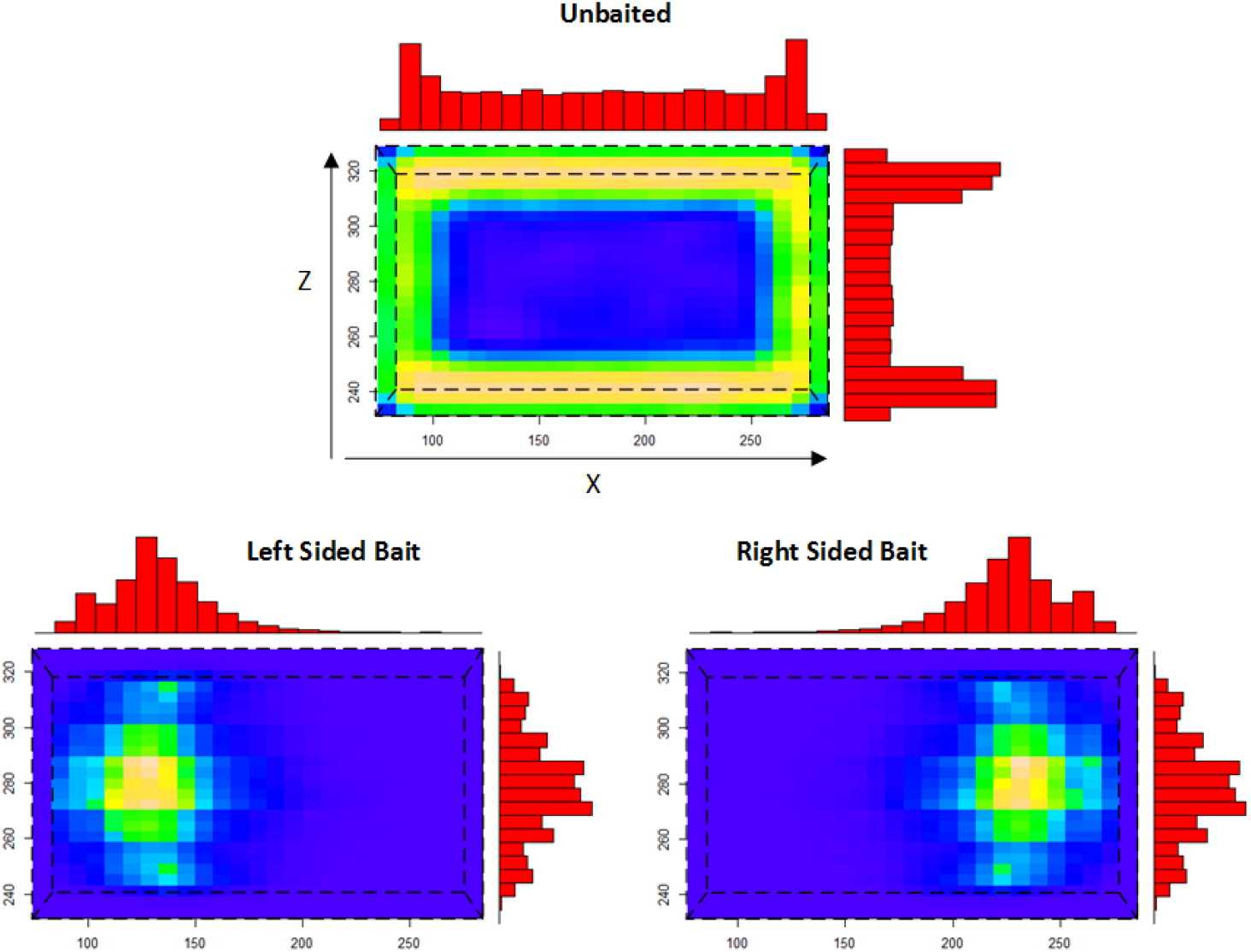
Distribution of bed net contact sites (shown from above, net shape indicated by dashed line). Spatial 2D heat map distribution of occupancy in the bed net XZ plane and 1D frequency distributions. There is relatively little contact at the top of the net in the unbaited condition where most contact occurs at the side net surfaces. In the baited conditions there are a large number of contacts on the top net surface at locations corresponding to the upper torso and head of the host. These contact sites correlate with bait orientation.

Under simulated human bait conditions, contact events showed a predilection for the top surface of the net and The X coordinates of the contact patterns were strongly influenced by the orientation of the human bait (Fig. 6, bottom). An illustration comparing the distribution of net contact numbers and spatial distribution patterns in unbaited and baited experiments is shown in Fig. 7. It has been suggested that strong air currents may affect host-seeking behavior and, potentially, subsequent mosquito distribution on bed nets [16]. We assessed the effect of noise on attractant dispersal, virtual mosquito host-seeking and the number and spatial distribution of bed net contact locations. Fig. 8 demonstrates the spatial effects showing, at low noise levels, a wider region of contact on the net surface (Fig. 8,A-E). At higher noise levels, the mosquitoes increasingly contact the net at the sides, as opposed to the top surface (Fig. 8, F-H). This change of net contact height with increasing noise (Fig. 9,A) is accompanied by a lower number of total net contacts (Fig. 9,B).

**Fig 7.**
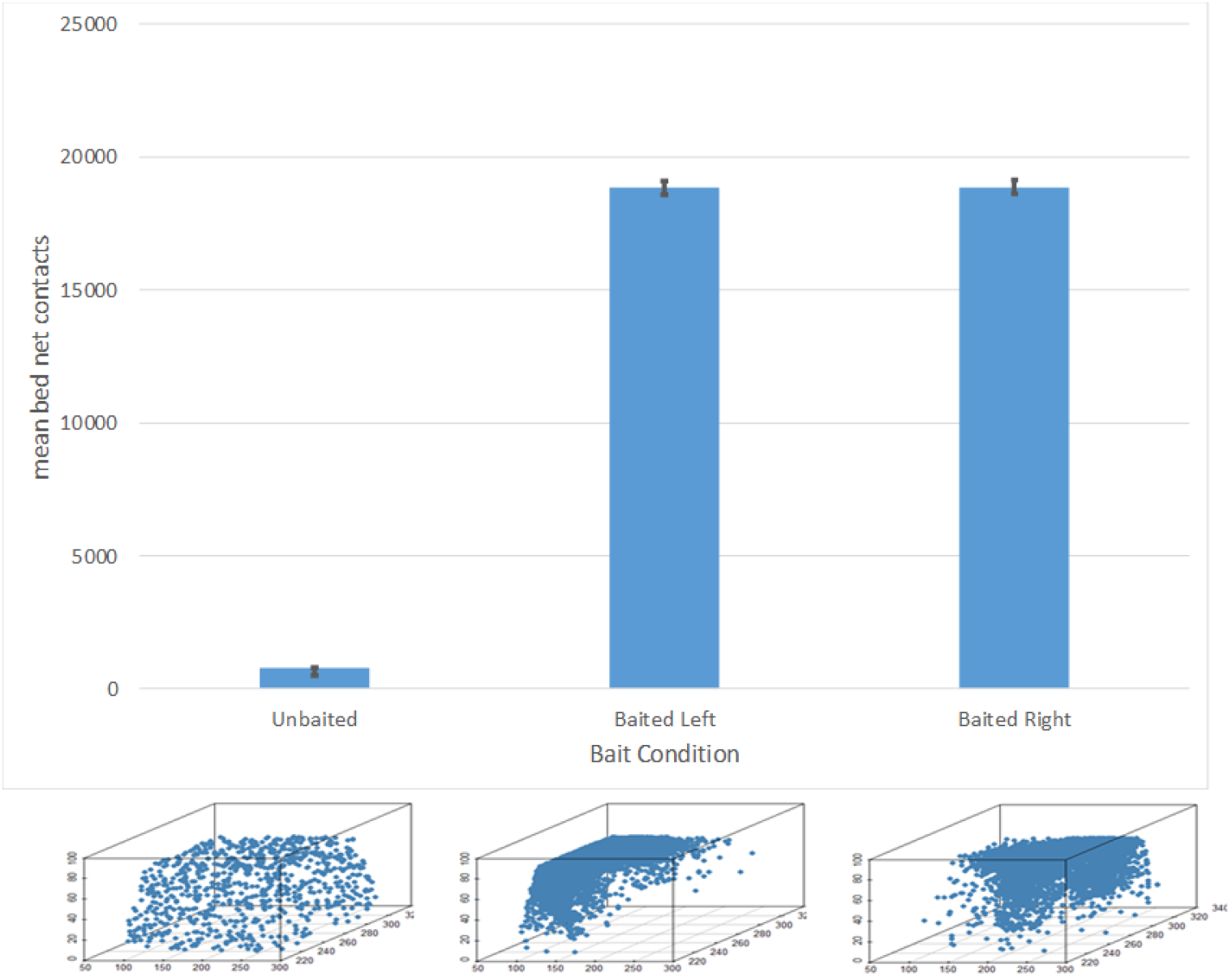
Mosquito net contact distributions. Top: Mean number of bed net contact events for each bait condition. Bottom: Example distribution of surface contacts in an experimental run. 3d scatter plot shows circles indicating individual contact sites on all surfaces of the 3d net. Unbaited (left) shows isotropic distribution of contacts. Left sided (middle) and right sided (right) bait conditions show distribution of contact sites over the head and torso of the bait stimulus location.

**Fig 8.**
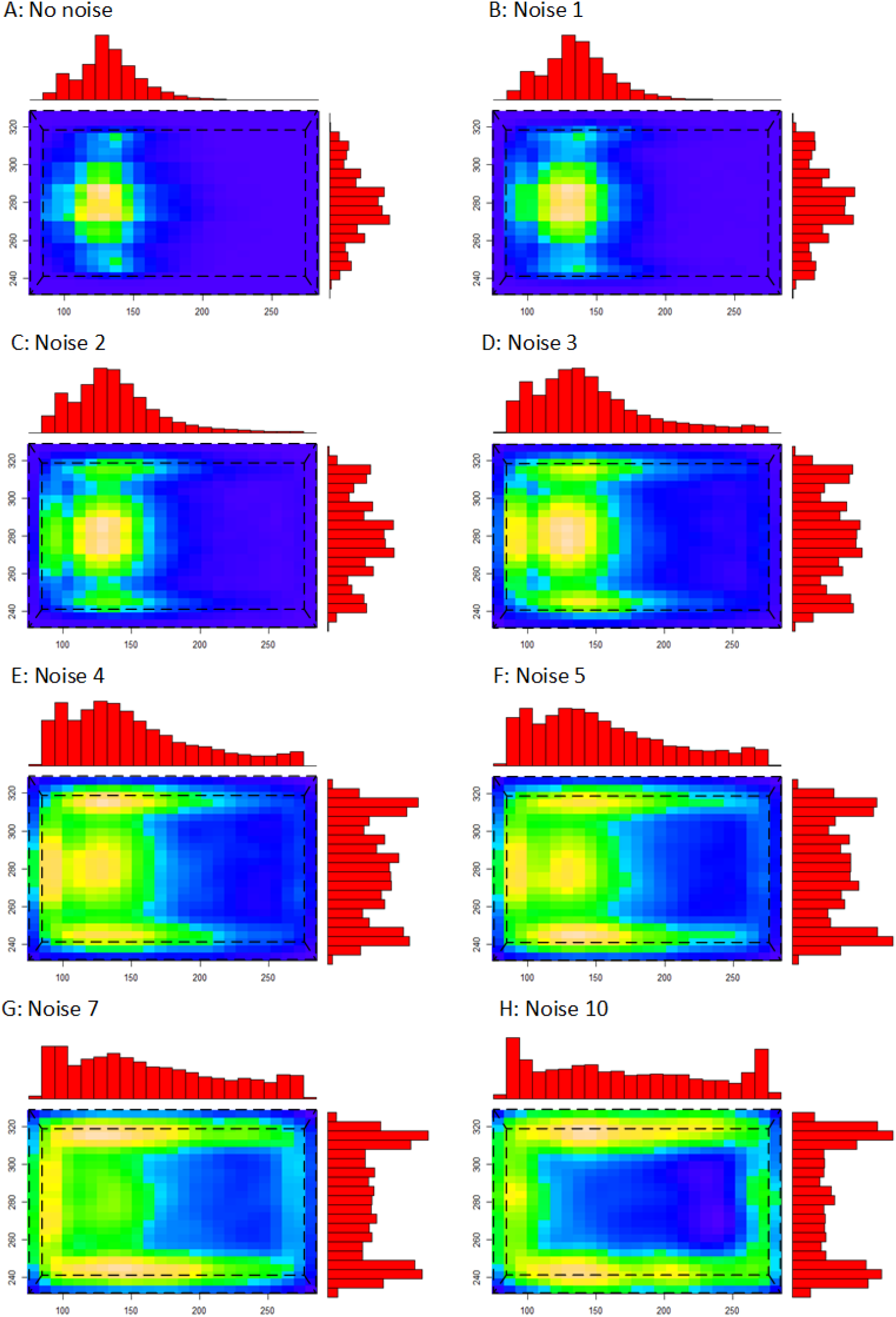
Effect of noise on attractant dispersal and mosquito bed net contact locations. Spatial heat map distribution of mosquito activity in the XZ plane (i.e. looking from above, net shape indicated by dashed line) under noisy dispersal conditions with a left-oriented host. A-F: noise multiplier from no noise contamination (A) up to 10 (B-F). Increasing noise spreads mosquito distribution across the top surface of the net and at higher noise levels, increases net contact at the sides of the net.

**Fig 9.**
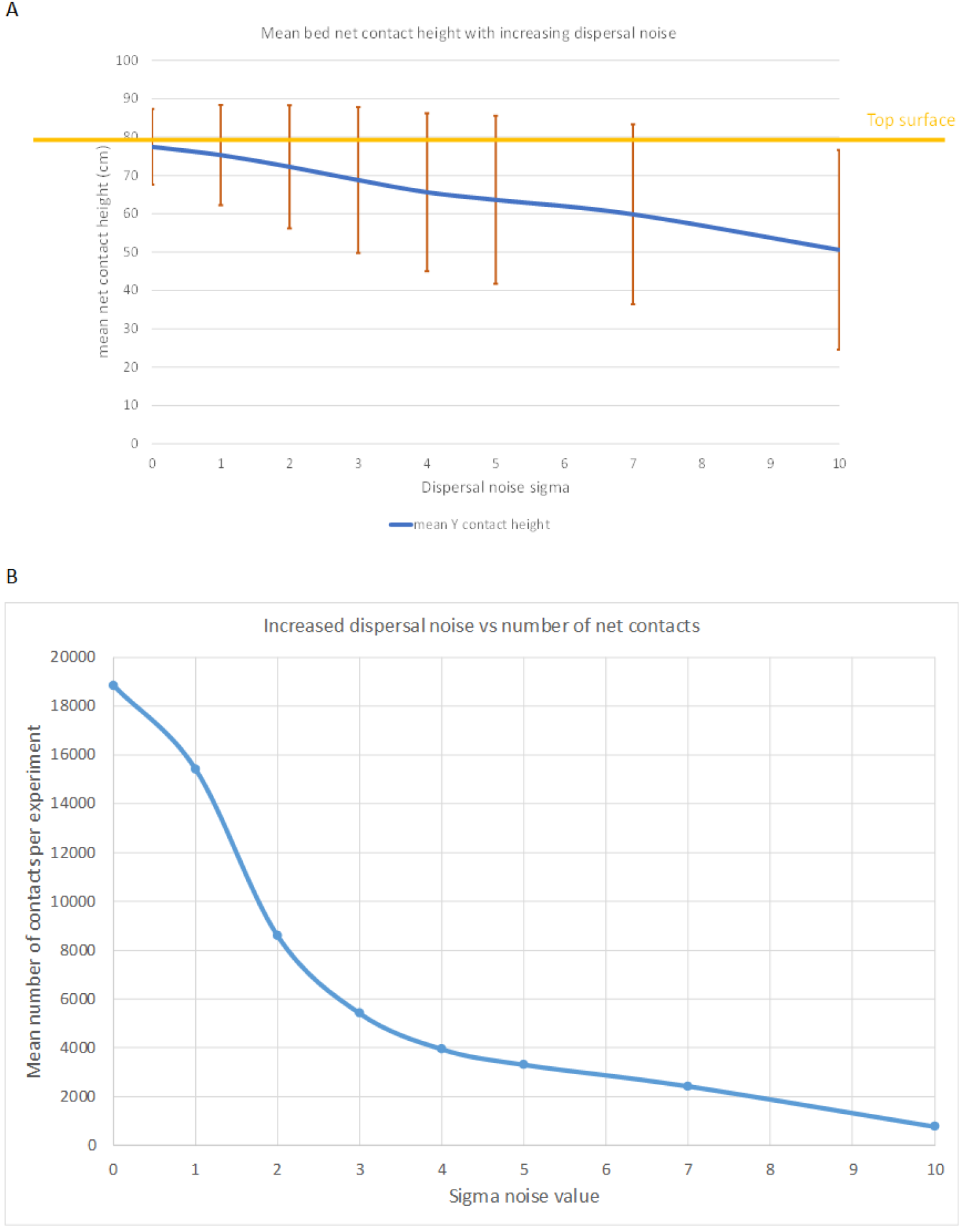
Effect of noise on total number of net contacts and contact height. A) mean net contact height decreases as dispersal noise increases, moving from the top surface (orange) to the sides of the net. B) The mean number of total net contacts decreases as dispersal noise increases.

### Peripheral bed net region occupancy

To assess peripheral occupancy of the regions surrounding the bed net we subdivided the recording volume into polyhedral shapes similar to, but not exactly identical, to the polygonal regions used in [10]. The reason for the differences is due to the 2D nature of the projection of the image tracking method and the slight difference in net orientation used in [10] (the net was tilted to allow better visualization of the top surface of the net for the capture cameras). The outer regions of the bed net in the model are divided into 18 regions which allows differentiation between the top surface of the net (12 sub regions), the two end edges of the net (edges corresponding to those near the head and feet) and the two sides of the net (4 regions consisting of arms and legs for both sides of the net). A 3D visualization of the peripheral regions is shown in S4. Occupancy of the regions is recorded during experimental runs. Each time a mosquito is located within a particular region, a counter for that region is incremented. The total mean region distribution data for control and baited conditions is summarized in S5.

A visual summary of flight tracks and peripheral region occupancy of the model output compared to experimental data in response to unbaited, untreated and insecticide treated nets is shown in Fig. 10 and supplementary data in S5. In the absence of a human host, there is slightly greater activity in regions to the side and ends of the bed net than in regions just above the net. This is partly due to the elongated nature of the virtual insectary but also because the two sides of the net present a larger surface area than the roof of the net. When a host is present in the net, however, the activity profile changes markedly. The attractant stimuli diffusing from the human host profile strongly attracts the mosquito population to the top regions surrounding the net. The presence of a baited net also affects flight activity in the peripheral regions in total, resulting in approximately 8 times as much peri-bed net activity compared to an unbaited net. A baited net coated with insecticide (0.1 ‘dose’ accumulation per contact, decrementing an initial health value of 100) exhibits the same strong attraction to the top surface of the net. Repeated contact, however, diminished the health of the mosquitoes until the population was entirely killed in a mean time of 36.34m (s.d. 2.56m).

**Fig 10.**
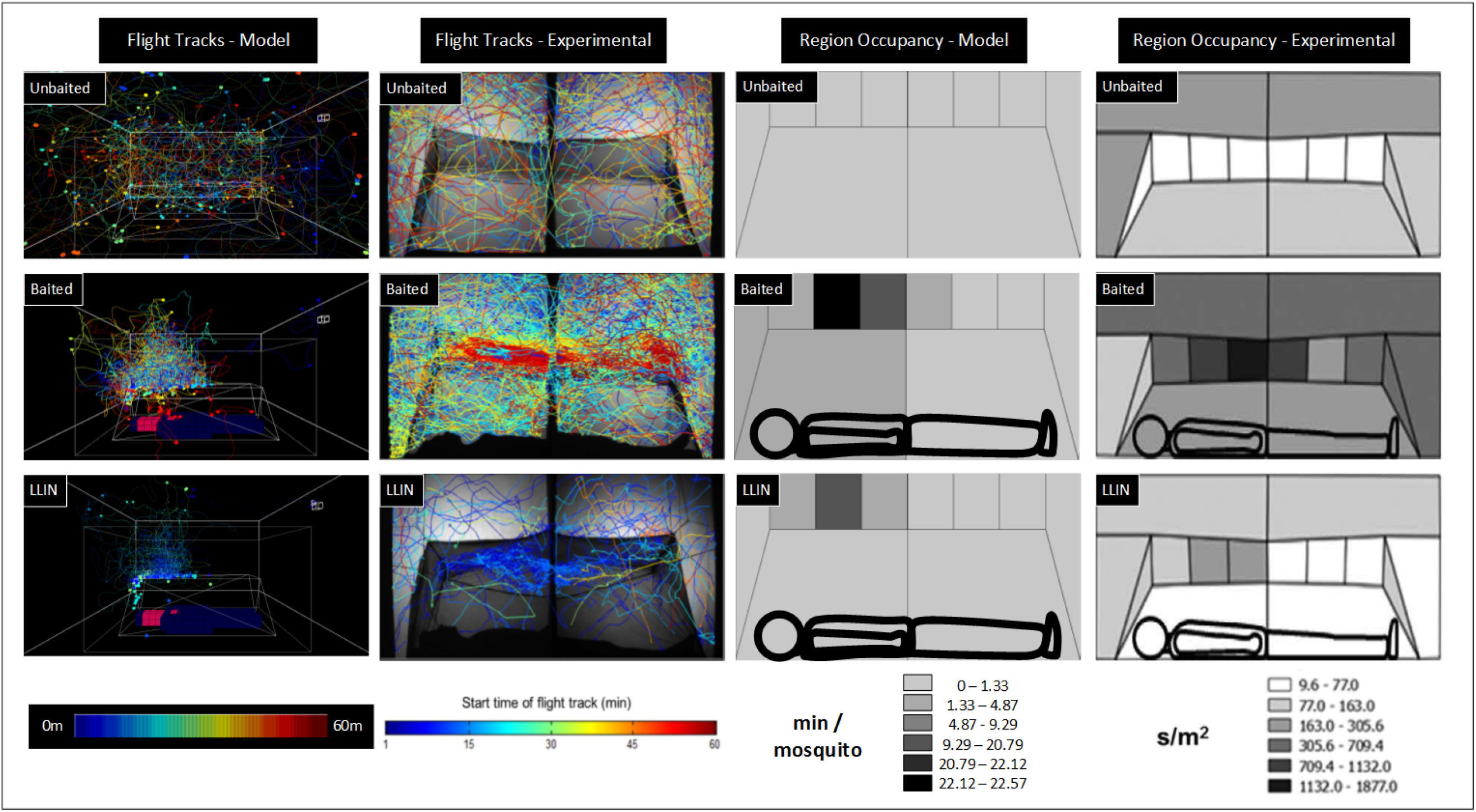
Spatio-temporal flight tracks and peri-bednet regional occupancy. Simulation (Cols 1 & 3) and experimental (Col 2 & 4) data. Track color indicates time (scale for flight tracks indicated below Cols 1 & 2). Region color indicates mean occupancy time indicated below Cols 3 & 4. Conditions shown are for an unbaited net (top row), an untreated net (middle row) and LLIN treated net (bottom row). Model tracks show a single mosquito for clarity and circular regions denote landing sites. Experimental images reproduced courtesy of [10].

## Discussion

The human home is exploited by numerous parasitic arthropods, including vectors of malaria, dengue, yellow fever, lymphatic filariasis, leishmaniasis and Chagas disease. In Africa, most cases of malaria are transmitted indoors, despite the wide range of behavioral preferences shown by those *Anopheles spp.* that are vectors [37], [38]. In turn, humans have exploited this behavior using a number of methods to target them at various locations inside the home. Of the methods used to date, insecticide-treated bed nets (ITNs/ LLINs) have been shown to be highly effective [1], and they remain a fundamental element in malaria prevention in Africa. As a first stage in developing models that capture vector behavior from house entry to exit, incorporating spatial movement and resting site preferences, this agent-based model of mosquito flight and host-seeking dynamics was developed, based on the actual conditions and their impacts on vector behavior measure previously.

To our knowledge, this is the first fine-grained model to simulate mosquito flight and distribution patterns in 3D space from quantification of actual behaviors recorded in earlier studies ([10], [28], [29]). Notably here, the patterns of net contact and flight in the peri-bed net region were accurately reproduced by the model.

It is the combined effect on this behavior of the attractants emanating from the host within the net and the potentially repellent or irritant properties of the insecticide treatment on the net, that determine whether or not a mosquito makes sufficient net contact to acquire a lethal dose. That this is accurately represented in the model is shown by figure 10. Without a host acting as attractant bait, activity around the bed net was relatively uniform (10, top row), reflecting the shape of the arena and positioning of the bed net. When a human host was present, the behavior of the mosquito population changed in response to different stimulus conditions, aggregating preferentially on the roof of the bed net (10, middle row). The foraging behavior of the virtual population measured by an occupancy metric based on the fraction of the arena explored, shifted from an ‘exploration’ behavior (wide spatial foraging within the arena) to an ‘exploitation’ behavior (a narrower spatial exploration as the population attempted to penetrate the net to take a blood meal). This difference in behavior was entirely provoked by the diffusion of the attractant plume from a host. With an untreated net this foraging and attempted penetration of the net persists over the entire hour whereas LLIN treated nets, in experimental and simulation conditions, dramatically reduced the activity of the mosquito population (10,bottom row).

We approximated turbulent dispersal of the attractant field using a noise parameter which added Gaussian noise during the diffusion method. Increasing dispersal noise resulted in fewer net contacts, a wider distribution patter and, at high noise values, a change in contact distribution from the top surface of the net to the sides of the net.

### Limitations of the model

These results demonstrate that a fine-grained modelling approach has utility as an in-silico method of performing virtual mosquito bio-assay experiments to explore 3D indoor flight behavior. As with all modelling approaches, limiting assumptions have been made. Limitations of the approach include the simplified and generic nature of the attractant stimulus (for example, visual stimuli are not represented). This spatially dispersed stimulus was chosen as it is the most well studied and because integration of stimuli from other sensory modalities are extremely complex [35], [39] and are yet to be fully understood. We acknowledge and stress that the mechanism of attractant dispersion by cellular automata-based methods is a simplification, partly due to previously noted debate regarding the relative contribution of difference stimulus types, but also necessitated by the desire for real-time simulation performance. Alternate methods of attractant dispersion and sampling (for example, computed by computational fluid dynamics methods) are a future possibility, although this would likely bias the model to particular avenues of stimulus dispersion. The model mosquitoes do not incorporate any approximation of energy expenditure and therefore do not display a tail-off in flight activity over time, as reported in [10]. Furthermore, the relative simplicity of the mosquito behavioral stimulus-response transitions was chosen to attempt to reduce the wide range of potential parametric influences on model behavior, a known issue with agent-based approaches. Nevertheless, despite these limiting assumptions, the model has demonstrated that it is able to reproduce the findings in [28], [29], most notably that the distribution of mosquito landing sites occurs predominantly on the top surface of the net and is affected by disturbances in dispersion by shifting this distribution to lower regions of the net. Furthermore the model also reproduces experimental tracking findings of [10], exhibiting similar flight tortuosity, peri-bed net distribution and activity decay in response to simulated insecticide treated bed nets.

### Application of the model and scope for further work

By expanding the parameter range, the model can accommodate far more than described here. For example, modifications might include vectors with different arrival patterns at the bed net, different host preferences, or the influence of a second human host in the room, or of a residual insecticidal treatment on the wall. Recently, we used an expanded version of the model to compare seven barrier bed net designs, which allowed us to rapidly identify the most promising candidate(s) to fast track for further evaluation [40].

## Supporting information

**S1 Simulated flight behavior in unbaited condition**. Video recording of simulated mosquito flight behavior in response to an unbaited bed net. Population of 25 mosquitoes.

**S2 Simulated flight behavior in baited condition**. Video recording of simulated mosquito flight behavior in response to a human occupied bed net. Population of 25 mosquitoes.

**S3 Schematic flowchart of mosquito behavior transition function.**

**S4 Subdivision of peri-bed net regions**. Subdivision of space peripheral to the bed net into 18 regions comprised of 3D polyhedra covering the top surface (coded as 0 — 11 white sub-regions, 0 — 5 (top strip), 6 — 11 (bottom strip)), short ends (12 and 13, (red and green respectively)) and side regions (14 — 17, blue (top) and magenta (bottom) respectively).

**S5 Peri-bed net region occupancy**. Mean time per mosquito (s) occupying each region surrounding the bed net for each condition. Regions and their sub regions indicated on left side. Mean counts and Standard Deviation per condition. Summary table indicating total activity per mosquito (m) within peri-bed net regions.

**S6 Evaluation of the effect of SA (sensor angle) and RA (rotation angle) parameters**. Chart illustrating how both parameters affect arena occupancy and path tortuosity and a description of how a continuous tortuosity metric was calculated.

## Acknowledgments

The authors gratefully acknowledge the support of the UK MRC Confidence in Concept funding and support from the Bill and Melinda Gates Foundation (opportunity number OPP1159078).

