## SupplementaryMaterial for "A minimal 3D model of mosquito flight behavior around the human baited bed net"

**Supporting information:** Contents are listed below and can on the following pages.

**S1 Video.**

Simulated flight behaviour in unbaited condition. Video recording of simulated mosquito flight behaviour in response to an unbaited bed net. Population of 25 mosquitoes.

**S2 Video.**

Simulated flight behaviour in baited condition. Video recording of simulated mosquito flight behaviour in response to a human occupied bed net. Population of 25 mosquitoes.

**S3 Fig.**

Schematic flowchart of mosquito behaviour transition function.

**S4 Fig.**

Subdivision of peri-bed net regions. Subdivision of space peripheral to the bed net into 18 regions comprised of 3D polyhedra covering the top surface (coded as 0 − 11 white sub-regions, 0 − 5 (top strip), 6 − 11 (bottom strip)), short ends (12 and 13, (red and green respectively)) and side regions (14 − 17, blue (top) and magenta (bottom) respectively).

**S5 Table.**

Peri-bed net region occupancy. Mean time per mosquito (s) occupying each region surrounding the bed net for each condition. Regions and their sub regions indicated on left side. Mean counts and Standard Deviation per condition. Summary table indicating total activity per mosquito (m) within peri-bed net regions.

**S6 Fig.**

Evaluation of the effect of SA (sensor angle) and RA (rotation angle) parameters. Chart illustrating how both parameters affect arena occupancy and path tortuosity and a description of how a continuous tortuosity metric was calculated.

**S1 Video.**

Simulated flight behaviour in unbaited condition. Video recording of simulated mosquito flight behaviour in response to an unbaited bed net. Population of 25 mosquitoes.

See attached video file
or follow link: https://youtu.be/VjdkkSMZiPU

**S2 Video.**

Simulated flight behaviour in baited condition. Video recording of simulated mosquito flight behaviour in response to a human occupied bed net.  Population of 25 mosquitoes.

See attached video file
or follow link: https://youtu.be/PZlOZjTkhuk

**S3 Fig.**

Schematic flowchart of mosquito behaviour transition function.

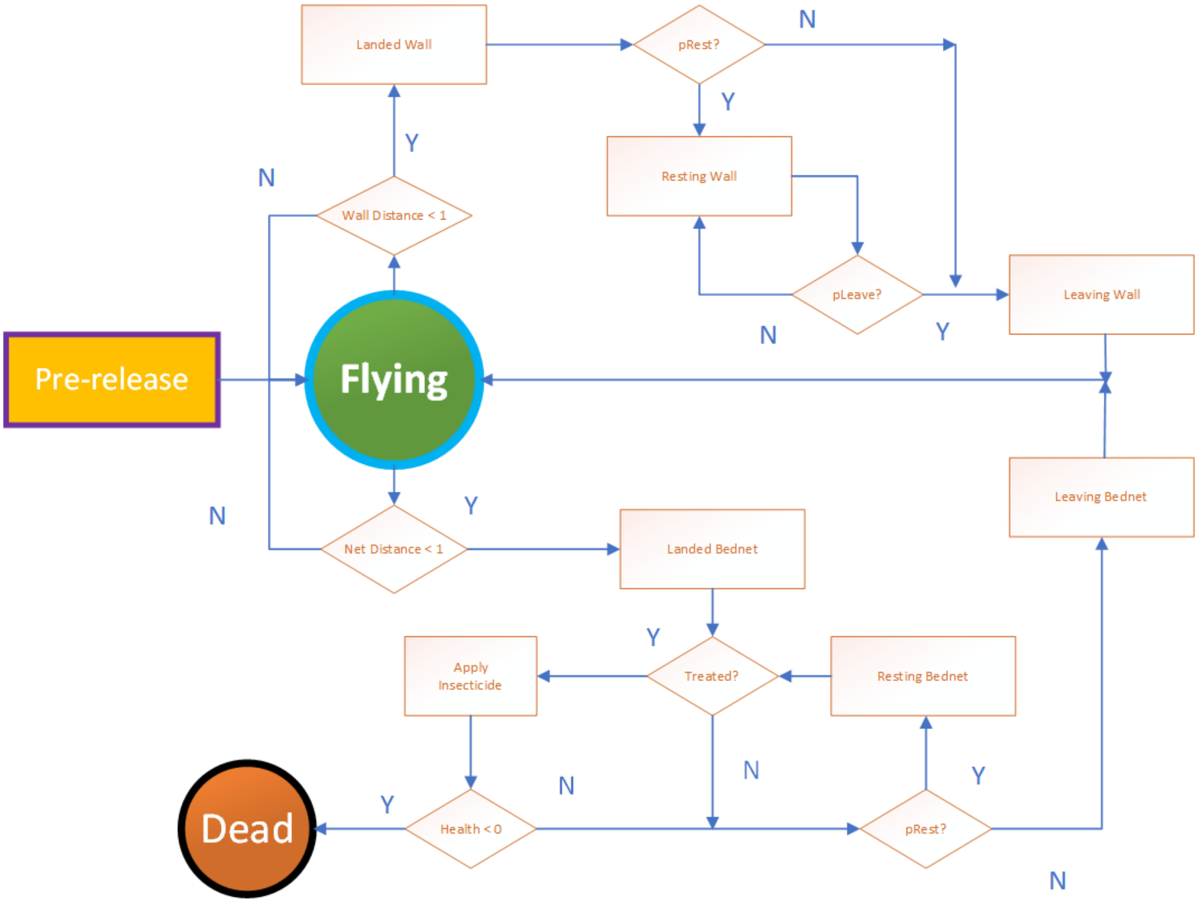

**S4 Fig.**

Subdivision of peri-bed net regions. Subdivision of space peripheral to  the bed net into 18 regions comprised of 3D polyhedra covering the top surface (coded  as 0 – 11 white sub-regions, 0 – 5 (top strip), 6 – 11 (bottom strip)), short ends (12 and 13, (red and green respectively)) and side regions (14 – 17, blue (top) and magenta (bottom) respectively).

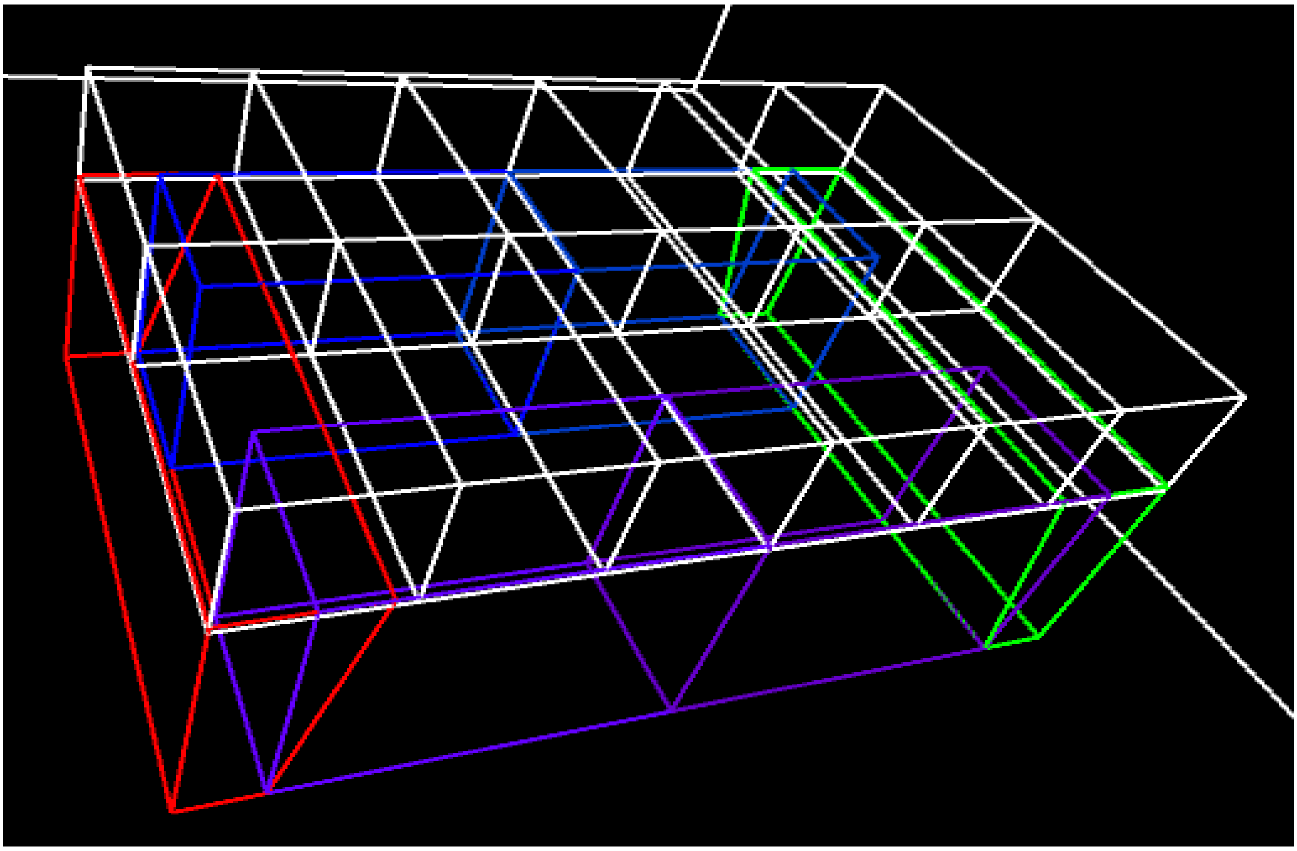

**S5 Peri-bed net occupancy Tables.**

Peri-bed net region occupancy. Mean time (s) per mosquito occupying each region surrounding the bed net for each condition. Regions and their sub regions indicated on left side. Mean counts and Standard Deviation per condition.

| Region | Sub Region | No Net or Bait 5 runs (s) | SD | Unbaited Net 20 runs (s) | SD | Left Sided Bait 20 runs (s) | SD | Right Sided Bait 20 runs (s) | SD | LLIN Treated Net 0.1/contact (s) | SD |
|---|---|---|---|---|---|---|---|---|---|---|---|
| Top | R0 | 10.49 | 0.82 | 10.57 | 0.99 | 95.46 | 2.51 | 0.58 | 0.22 | 42.83 | 3.71 |
| | R1 | 10.22 | 0.55 | 10.34 | 0.91 | 709.85 | 9.81 | 6.01 | 1.18 | 316.30 | 18.10 |
| | R2 | 10.73 | 0.44 | 10.41 | 0.97 | 347.15 | 6.63 | 46.78 | 2.64 | 153.24 | 11.50 |
| | R3 | 10.76 | 0.31 | 10.51 | 0.98 | 49.93 | 2.57 | 348.43 | 6.57 | 22.78 | 2.82 |
| | R4 | 10.21 | 0.75 | 10.38 | 1.25 | 7.09 | 1.17 | 689.45 | 10.39 | 3.44 | 0.92 |
| | R5 | 10.05 | 1.12 | 10.57 | 0.91 | 0.92 | 0.32 | 96.33 | 2.96 | 0.50 | 0.32 |
| | R6 | 9.81 | 0.45 | 10.31 | 0.88 | 93.98 | 3.24 | 0.71 | 0.31 | 41.98 | 3.46 |
| | R7 | 10.36 | 0.36 | 10.49 | 0.65 | 642.50 | 9.46 | 5.42 | 0.85 | 287.58 | 17.47 |
| | R8 | 10.05 | 0.76 | 10.52 | 0.74 | 272.47 | 6.82 | 39.08 | 2.07 | 119.71 | 9.72 |
| | R9 | 9.89 | 0.40 | 10.27 | 0.85 | 36.69 | 2.10 | 269.06 | 5.62 | 16.69 | 2.11 |
| | R10 | 10.70 | 0.97 | 10.17 | 0.93 | 5.11 | 0.82 | 664.58 | 8.01 | 2.49 | 0.96 |
| | R11 | 10.82 | 0.98 | 10.58 | 0.85 | 0.81 | 0.24 | 95.84 | 2.90 | 0.38 | 0.23 |
| Ends | R12 | 29.67 | 1.09 | 29.03 | 3.12 | 94.99 | 6.14 | 0.72 | 0.47 | 41.53 | 5.47 |
| | R13 | 30.29 | 1.29 | 29.54 | 2.03 | 0.77 | 0.53 | 96.15 | 5.88 | 0.40 | 0.42 |
| Sides | R14 | 27.36 | 0.91 | 26.64 | 2.90 | 81.24 | 4.89 | 3.97 | 0.76 | 36.04 | 4.34 |
| | R15 | 27.55 | 1.79 | 26.43 | 2.75 | 3.75 | 0.84 | 81.75 | 3.89 | 1.54 | 0.76 |
| | R16 | 26.17 | 1.34 | 25.98 | 2.70 | 72.54 | 4.62 | 3.38 | 0.88 | 33.73 | 5.49 |
| | R17 | 25.79 | 1.45 | 25.94 | 1.96 | 3.15 | 0.71 | 73.54 | 4.90 | 1.51 | 0.71 |
| **Total (s)** | **All** | **290.91** | | **2518.40** | | **2521.78** | | **2520.09** | | **1122.68** | |

Summary of the mean time spent (minutes per mosquito) in all regions surrounding the bed net. In the unbaited condition there is only 4.81 minutes spent in regions surrounding the net, compared to 42 minutes in the baited condition. The effect of a treated bed net reduces the regional occupancy to 18.71 as the mosquito population is killed off.

| | Total time spent in Regions Surrounding Bed Net (m) |
|---|---|
| No Net or Bait Present | 4.85 |
| Unbaited Net | 4.81 |
| Untreated Net | 42.00 |
| LLIN Treated Net | 18.71 |

**S6 Fig.**

Evaluation of the effect of SA (sensor angle) and RA (rotation angle) parameters. Chart illustrating how both parameters affect arena occupancy and path tortuosity and a description of how a continuous tortuosity metric was calculated.

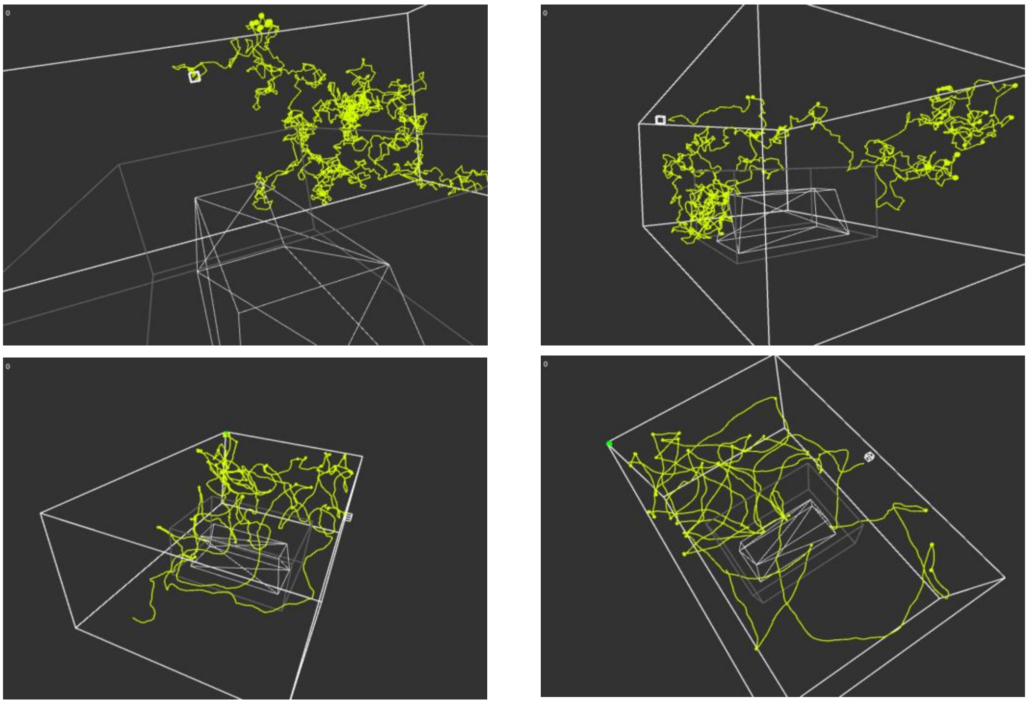

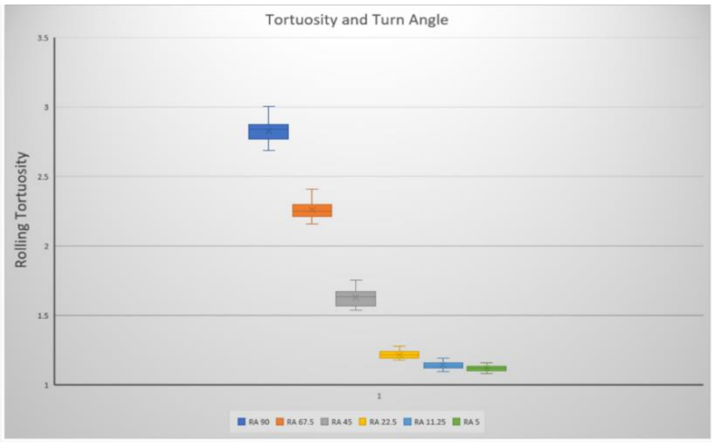


Virtual mosquito turn angle parameter affects foraging and occupancy behavior and flight path tortuosity. Top: Examples of 3D flight paths from a single agent in unbaited arena at turn angles of $90^0$ (top left), $67.5^0$ (top right), $22.5^0$ (middle left) and $11.25^0$ (middle right). Bottom: Plot of rolling tortuosity metric at different turn angles. Rolling tortuosity is computed per mosquito, per time step as a comparison between 50 contiguous flight movements. The window covers all 50 scheduler steps of uninterrupted movement throughout the entire 1hr experiment and path comparison is by comparing mosquito positional vector at each of these 50 steps against the Euclidean straight line between start point $_t0$ and end point $_t50$. A value of 1 would indicate a perfectly straight track between the points. Values greater than 1 indicate higher path tortuosity.
